# Lossless compression of protein databases for efficient and accurate metagenomic sequence classification with Centrifuger

**DOI:** 10.64898/2026.09.17.752512

**Authors:** Li Song

## Abstract

We present a lossless compression algorithm for indexing a protein database while supporting fast taxonomic classification in the method Centrifuger. The algorithm is a new scheme of the previously proposed run-block compression algorithm to reduce the size of the Ferragina-Manzini (FM) index, and it scales better with alphabet size than the original. On the RefSeq prokaryotic and viral protein sequences, Centrifuger reduces the memory footprint by over a third compared to the method Kaiju that builds on a plain FM-index, while having comparable running time. Furthermore, the compressed FM-index is lossless and can locate matches of arbitrary length, which helps Centrifuger achieve greater accuracy than Kraken2, a k-mer-based taxonomic classification method. We leverage the computational efficiency of Centrifuger to create an index of size 182 GB for classifying the reads against the full nr database that contains about 250 billion amino acid characters. Using this index, Centrifuger reveals different SARS-CoV-2 infection states and viral transcriptome profiles across human cell types from single-cell RNA-seq data.

## Introduction

Metagenomic sequencing has become a routine technique to investigate the microbiome composition in environmental^1,2^ and human disease^3^ samples. Taxonomic classification is the task of finding the taxonomy ID for each read in the data, which can be utilized to infer the microbiome abundances^4^ and detect low-abundance taxa^5^. Therefore, it has become a crucial step in metagenomic data analysis. The common approach for taxonomic classification is to search read patterns against the comprehensive microbial genome database, such as RefSeq^6^, GenBank^7^, or GTDB^8^. Despite the rapidly increasing volume of the database, there are still many underrepresented species and uncharacterized variants, and reads from such genomes are often unclassifiable due to large sequence divergence. Since protein sequences are relatively conserved in nature, taxonomic classification against a protein database can recover relevant taxonomic information for those underrepresented species.

Indexing the genome or protein database with k-mers is the most common paradigm for fast taxonomic classification methods, such as Kraken2^9^, CLARK^10^, KMCP^11^, Ganon2^12^, and Metabuli^13^. When the database grows large, a fixed-size k-mer can be found in distantly related species by chance, and loses classification specificity, i.e., the capability to classify at the lower taxonomic ranks like the species level^14,15^. This issue is exacerbated when these classifiers discard sequence information, like by using minimizers, discriminative k-mers, and probabilistic data structures, to reduce the index size. The Ferragina-Manzini (FM)^16^ index, which builds on the Burrows-Wheeler transform (BWT), is a full-text indexing paradigm. It is relatively memory efficient and supports finding matches of arbitrary length.

Therefore, the FM-index can address the aforementioned issues of k-mer indexing. The methods Centrifuge^17^ and Kaiju^18^ have pioneered efficient taxonomic classifications using the FM-index against the genome database and protein database, respectively.

However, the size of the plain FM-index increases linearly with the database size and can become prohibitive for ever-growing database volumes. For example, the pre-built Kaiju index for the subset of 2024-08-25 NCBI nr database containing only archaea, bacteria, and virus protein sequences already needs 219 GB of memory according to Kaiju’s website. The common databases for taxonomic classification, such as RefSeq, contain a mild level of repetitiveness, which is reflected as run structures, i.e., a stretch of substring with the same character, in the BWT sequence. In our previous work, we designed a compression scheme, called run-block compressed BWT (RBBWT)^19^. It is more effective in compressing and indexing genome databases with mild redundancy than data structures like Lempel-Ziv family indexes^20^, context-free grammars^21^, r-index^22^ and move structure^23^ that are excellent for pangenomes. When testing RBBWT on nucleotide and amino acid (AA) alphabets, we observed that RBBWT had a smaller efficiency advantage for the AA alphabet, especially when compared with the run-length compressed BWT (RLBWT)^24^, the data structure underlying r-index. Motivated by this, we propose a new variation of RBBWT, called run-block compressed BWT with one sequence (RBBWT-1S), which has a better time-space trade-off for a large alphabet size. We implement it in Centrifuger for taxonomic classification against the protein database. With RBBWT-1S, the Centrifuger index for the full 2026-01 nr database containing about 250 billion AAs is only of the size 182 GB. Centrifuger has options to parse information like the cell barcode from a read, so it is readily applicable to analyze microbial reads in single-cell RNA-seq (scRNA-seq) data. Since many RNA-seq reads come from protein-coding genes, searching the protein database also facilitates interpreting the microbial transcriptome profile in host cells.

## Results

### Method overview

Centrifuger is a taxonomic classification method that can find the taxonomy IDs for each read by searching against the protein database using the FM-index (**Figure 1**). To reduce the FM-index size for protein sequences, Centrifuger adopts a novel compression representation called RBBWT-1S. Compared with the previously proposed RBBWT representation, which splits the BWT into two sequences, RBBWT-1S compresses the BWT into one sequence with an auxiliary bit vector for each letter. RBBWT-1S is substantially faster than RBBWT for pattern search when the alphabet size is large; therefore, it is more suitable for representing the protein sequences. During the taxonomic classification, for a read strand, Centrifuger translates it into the three possible reading frames and greedily searches for semi-maximal matches for each frame. Specifically, it searches from the end of the translated AA sequence and extends the match backward until reaching a mismatch. Centrifuger then skips that mismatched AA and starts a new backward search from the position before the mismatch. Following an adaptation of Centrifuge’s empirical scoring function that sums the squares of the adjusted semi-maximal match lengths, Centrifuger selects the reading frame with the highest score. Unlike the standard FM-index that samples the suffix array elements, Centrifuger samples the sequence ID, e.g., the accession ID. This information is utilized to identify the sequence IDs for each match. The sequence IDs with the highest score can be converted to the taxonomy ID. In the case of too many sequence IDs with the highest score, Centrifuger will sweep their taxonomic common ancestors across the taxonomic ranks until the number of reported taxonomy IDs is within the user-specified threshold (default threshold 1).

**Figure 1.**
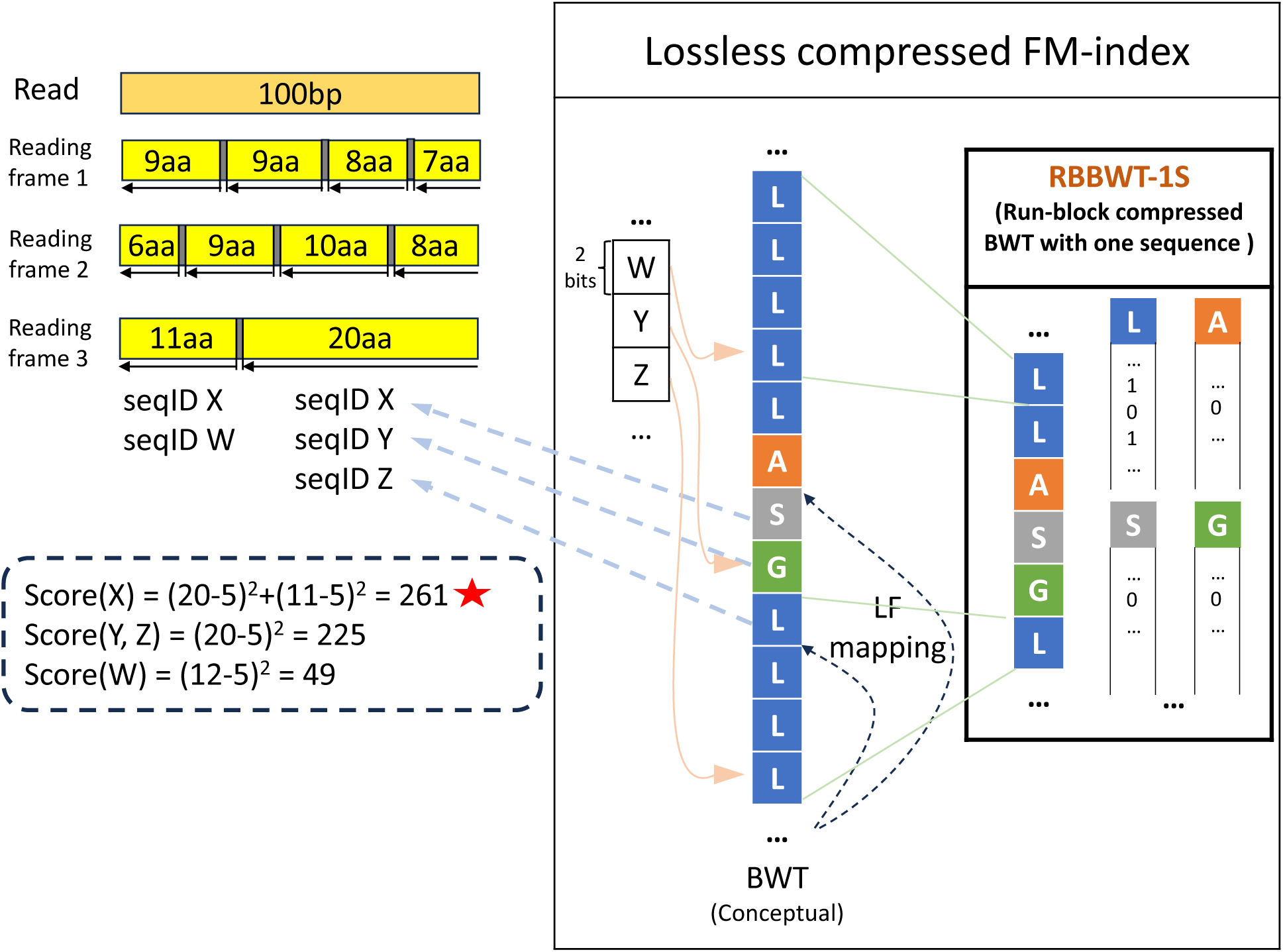
Overview of Centrifuger on taxonomic classification for protein sequences Left: An example of classifying the read forward strand. Centrifuger first translates the 100-bp read into three possible reading frames. For each reading frame, Centrifuger searches from its end and applies the backward search to extend the match until reaching a mismatch. For the third reading frame, this yields the first 20-AA exact match on three sequences {X, Y, Z} in the database. Centrifuger then skips the mismatch and restarts the search again, finding the second 11-AA match on two sequences {X, Y}. Centrifuger then scores each match and classifies the read to the highest-scoring sequences, where the example read is classified to the sequence X with the score 261. Right: Losslessly compressed FM-index with RBBWT-1S. In the example of compressing the BWT sequence “LLLLLASGLLLL”, RBBWT-1S represents it as the sequence “LLASGL” when the block size is 4, along with a bit vector for each letter. The sampled SA values in standard FM-index are converted to sequence IDs, which are shown as W, Y, Z in the example.

### The computational efficiency of run-block compression with one sequence

We compared the rank query speed for the wavelet tree, run-length compressed, run-block compressed and run-block-with-one-sequence compressed representations when the alphabet set size increased.

The rank query finds the number of occurrences of a given character up until a position in a sequence. This operation is essential in the LF mapping and backward search with the FM-index. We first created a BWT sequence from 282 genomes of the genus *Legionella*. The run structure of this BWT sequence served as the template with the average run length about 7.6. When generating a sequence with an alphabet size of *σ* = |Σ|, we randomly mapped the character in a run to a character from the alphabet set Σ={‘0’, ‘1’, …, ‘0’+*σ*-1}, where the addition was with respect to the ASCII code. For example, the procedure would create a sequence comprised of the characters ‘0’, ‘1’, ‘2’, and ‘3’ when setting *σ* as four. We also ensured that adjacent runs were mapped to different characters, so the run patterns remained the same. In this comparison, we observed that RBBWT-1S was faster than RBBWT, and the speed gap became larger when increasing alphabet size (**Figure 2A**). For the nucleotide alphabet, e.g., *σ* = 4, RBBWT-1S was only slightly faster than RBBWT, where the rank query took about 17.8 ns and 21.6 ns, respectively. When increasing *σ* to 21, which corresponded to 20 AA letters and the special end-of-sequence symbol ‘$’, the speed advantage of RBBWT-1S became clear. It was 3.2 times and 1.6 times as fast as RLBWT and RBBWT, respectively, and only took 45% more time than the standard wavelet tree representation.

**Figure 2.**
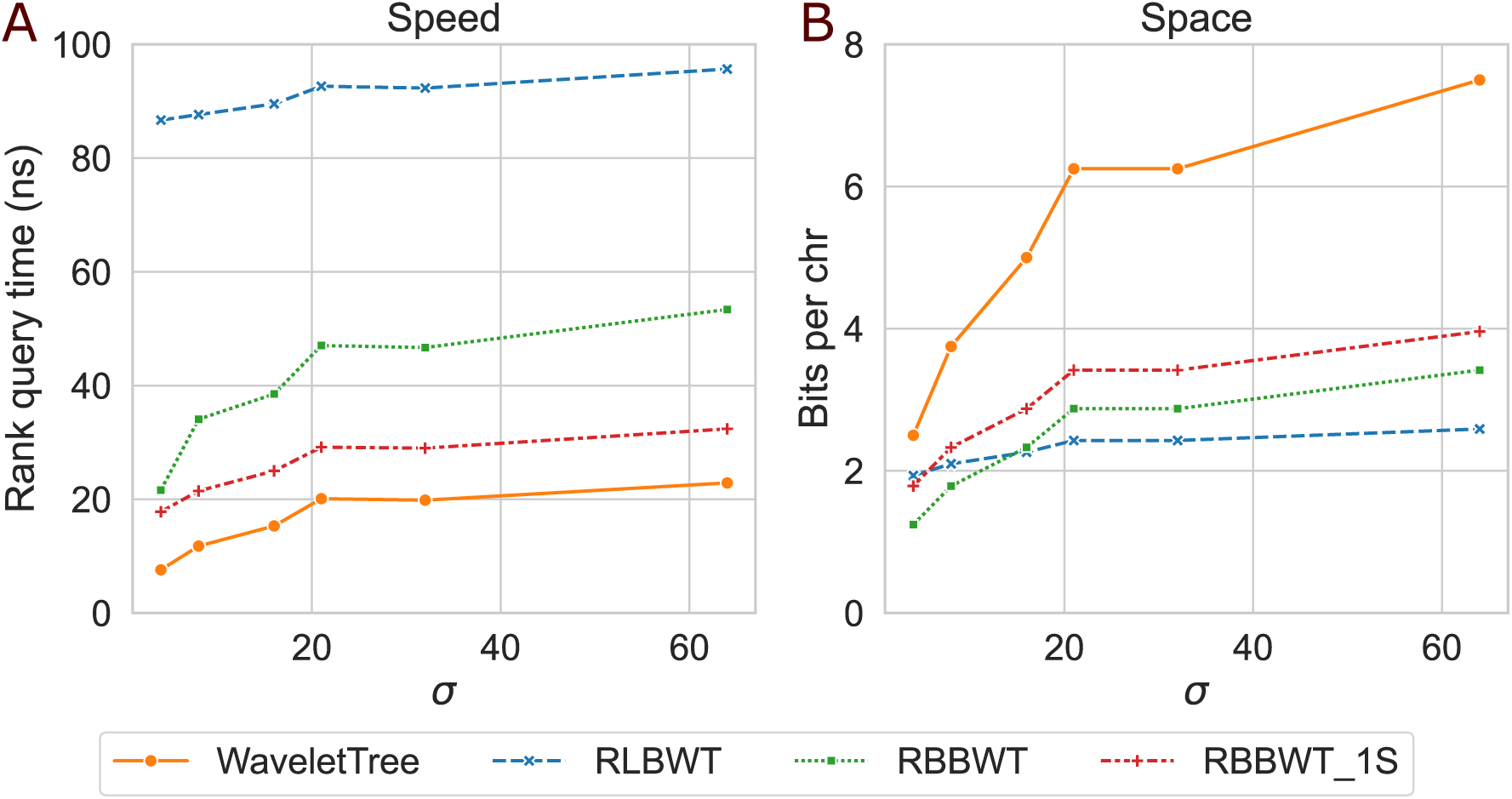
Computational efficiency of the wavelet tree, RLBWT, RBBWT, and RBBWT-1S. (A) Rank query time in ns (B) Space usage in bits per character

We next compared the space usage of these sequence representation data structures when varying the alphabet size. RBBWT-1S was less memory-efficient than other compression schemes (**Figure 2B**). For example, when *σ* was 21, RBBWT-1S needed 40.8% and 18.9% more space than RLBWT and RBBWT, respectively. Nevertheless, its compression ratio was still good, and it took 54.7% of a wavelet tree’s space. We observed that RLBWT compression was least sensitive to the alphabet size and became the most space-efficient representation when *σ* reached 16, suggesting that RLBWT could be suitable for a general alphabet set even when the average run length was small. In sum, RBBWT-1S achieved a good trade-off between time and space, especially for AA alphabet set.

### Performance on classifying simulated datasets

We compared Centrifuger with Kraken2 and Kaiju on 1 million 100-base-pair (bp) paired-end reads simulated by the method ART^25^ (**Figure 3A, Table S1**). The classifiers built their indices using the protein sequences from 67,498 prokaryotes (bacteria+archaea) and viruses with complete genomes in RefSeq. The genome sequences utilized to generate the simulated reads were downloaded at the same time as the protein database to maximize consistency. Classification accuracy was measured separately at different taxonomic ranks. We define a classification as true positive (TP) for a taxonomic rank if the classified taxonomy ID matches the ground truth or is in the ground truth’s taxonomic descendants. We defined the positive set (P) as reads for which a classifier reports a taxonomy ID, and the true set (T) as all the queried reads. Therefore, we could define the sensitivity as TP/T, corresponding to the fraction of correctly classified reads among all the queried reads. Precision was defined as TP/P, meaning the fraction of correctly classified reads with respect to classified reads. On this simulated dataset, Centrifuger was consistently more sensitive than Kaiju and Kraken2 with comparable precision. For example, at the genus level, Centrifuger was 6.2% and 4.1% more sensitive than Kaiju and Kraken2, respectively, and with the precision almost identical to Kaiju’s (0.993 for Centrifuger and 0.994 for Kaiju) and 1.4% higher than Kraken2’s.

**Figure 3.**
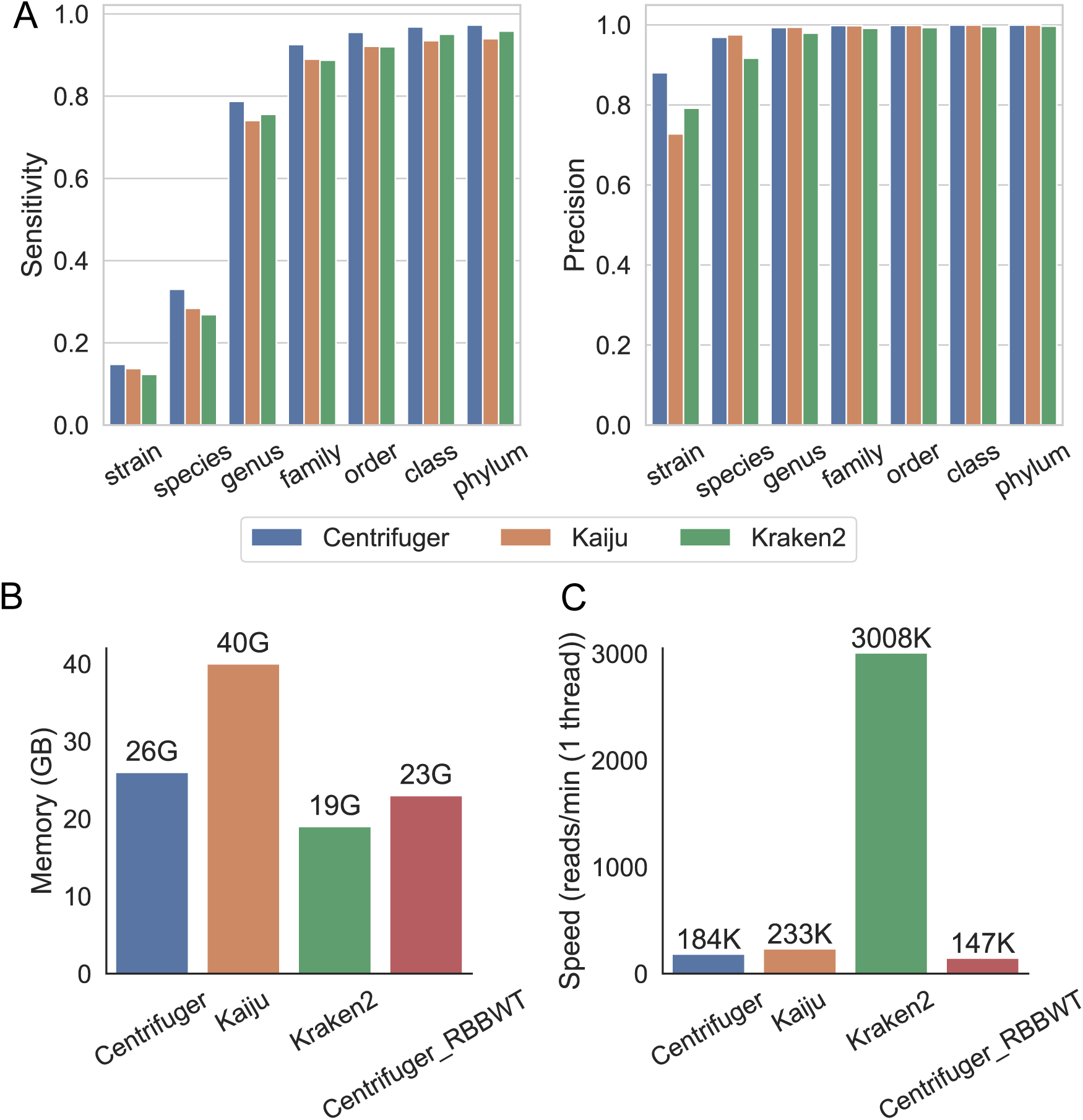
Performance of Centrifuger, Kaiju, and Kraken2 on the simulated data generated from August 2025 RefSeq prokaryotic and viral genomes (A) Sensitivity (left) and precision (right) of each classifier across taxonomic ranks. (B) Peak memory usage of each classifier and Centrifuger_RBBWT. (C) Single-thread classification speed of each classifier and Centrifuger_RBBWT

We examined the computational efficiency using this dataset. Centrifuger consumed 25 GB of memory for the protein database containing about 25 billion AAs (**Figure 3B**). Although Centrifuger took 38.9% more space than the lossy minimizer index of Kraken2, its space was about 65.8% of Kaiju’s, corresponding to the size of a plain FM-index. We note that the Centrifuger and Kaiju indexes stored sequence IDs in addition to taxonomy IDs, so they encoded more information than Kraken2. Despite searching on a compressed data structure, Centrifuger’s speed was comparable to Kaiju, where Centrifuger could process 184K reads per minute (reads/min) for one thread and was only 21.0% slower than Kaiju’s (**Figure 3C**). Although Kraken2 was ultrafast and could process over 3 million reads/min with a single thread, Centrifuger and Kaiju were still efficient. For example, they could process a sample with 50 million read pairs within one hour using multithreading, such as with 8 threads. We also tested a version of Centrifuger that used RBBWT as the underlying index compression scheme, denoted as Centrifuger_RBBWT (**Figure 3B,C**). Centrifuger with RBBWT-1S, the version proposed in this study, had 25.2% higher single-thread throughput than Centrifuger_RBBWT while consuming only 13.0% more memory. This confirmed the advantage of selecting RBBWT-1S over RBBWT for representing protein sequences in practice.

We further examined the performance of these methods when the ground-truth taxa were missing in the database (**Figure S1**). We created a protein database by keeping the protein sequences from one species per genus, and removed 53,385 previously simulated genomic reads from the representative species. The performance of Centrifuger and Kaiju was close. Their F1 scores, i.e., 2*sensitivity*precision/(sensitivity+precision), differed most at the species and genus levels, in which Centrifuger’s F1 scores were 5.9% and 1.7% higher than Kaiju’s, respectively. Centrifuger and Kaiju outperformed Kraken2 in sensitivity while having comparable precision. For example, Centrifuger’s sensitivity was 7.9% and 4.9% higher than Kraken2’s at the genus and family level, respectively.

We also benchmarked the three classifiers on 10 simulated samples from the strain-madness dataset of the Critical Assessment of Metagenome Interpretation 2 (CAMI2)^26^ study (**Figure 4**). The classifiers utilized the same index as in the ART-generated simulated data evaluation. Centrifuger achieved higher sensitivity, especially at the species and genus levels, than Kaiju and Kraken2, while having comparable or higher precision. For example, at the genus level, Centrifuger’s mean sensitivity was 19.6% and 29.3% higher than Kaiju’s and Kraken2’s, respectively, and the mean precision was 0.4% lower than Kaiju’s and 10.1% higher than Kraken2’s.

**Figure 4.**
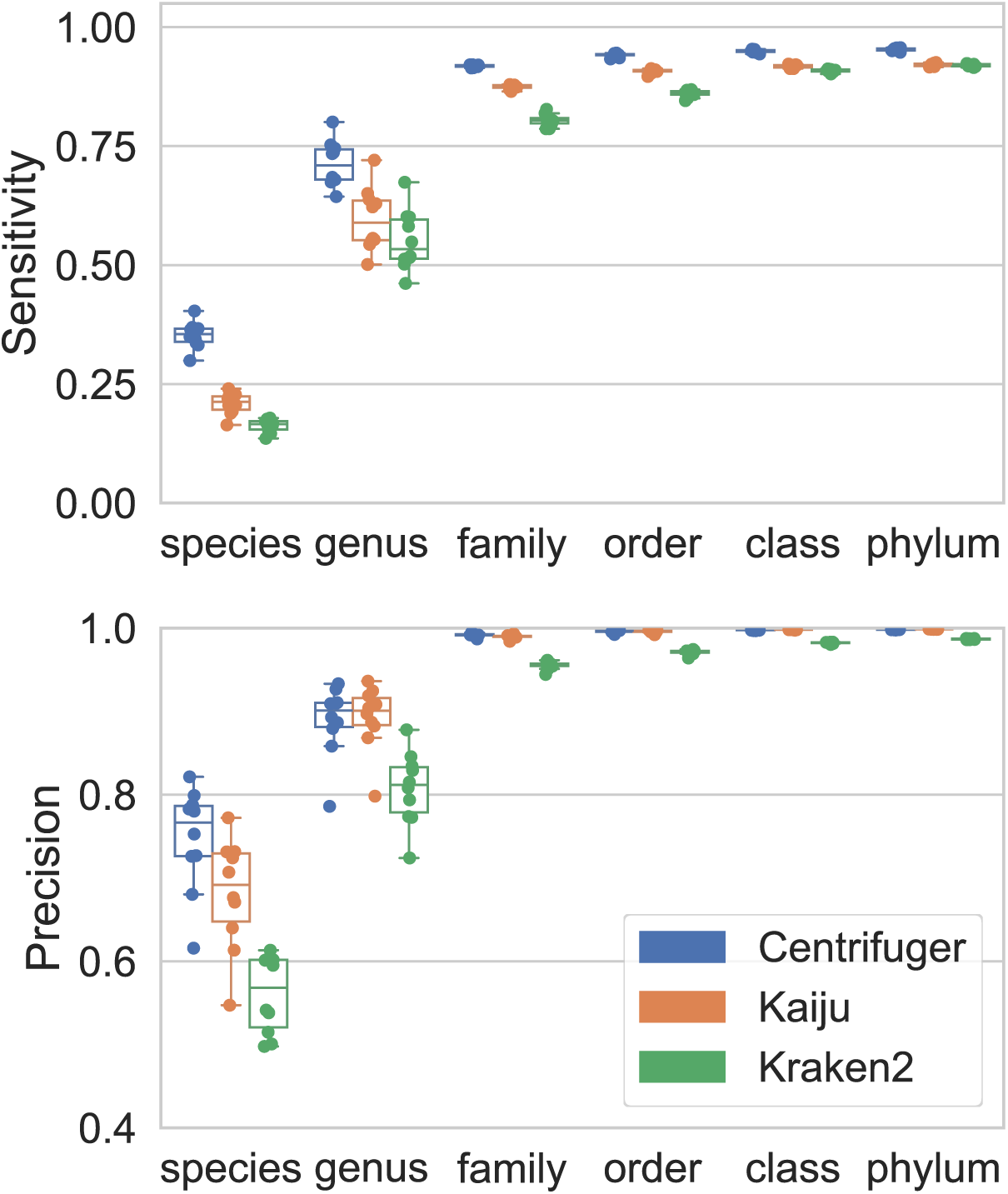
Sensitivity (top) and precision (bottom) of Centrifuger, Kaiju, and Kraken2 across taxonomic ranks on the 10 simulated datasets from CAMI2

### Performance on classifying bacteria whole-genome sequencing data

We examined Centrifuger, Kaiju, and Kraken2 on real datasets by utilizing 200 bacterial whole-genome sequencing (WGS) datasets. The protein database for the taxonomic classification was identical to the one used in the simulated data evaluation. For each WGS dataset, the true taxonomy ID was obtained from the NCBI SRA RunInfo. Among them, 100 datasets had the ground-truth species in the index (species-in), and 100 did not but had the ground-truth genus in the index (species-not-in). The accuracy of these classifiers was examined at the genus level. For the species-in scenario, Centrifuger was consistently more sensitive than the other methods, with the mean sensitivity 7.3% and 9.8% higher than Kaiju’s and Kraken2’s, respectively (**Figure 5A**). The three methods had comparable mean precision, where Centrifuger was 1.5% less precise than Kaiju but 2.1% more precise than Kraken2. In the evaluation, we also incorporated the results from Centrifuger searching against the genome database that was used to generate the simulated dataset. We denoted this mode as Centrifuger_genome. It achieved substantially higher mean sensitivity than protein-based classifications, while having similar mean precision, where its mean sensitivity was 6.3% higher than Centrifuger’s. This suggests that classifying against a protein database yielded less taxonomically specific results than against the genome database when the species was in the database.

**Figure 5.**
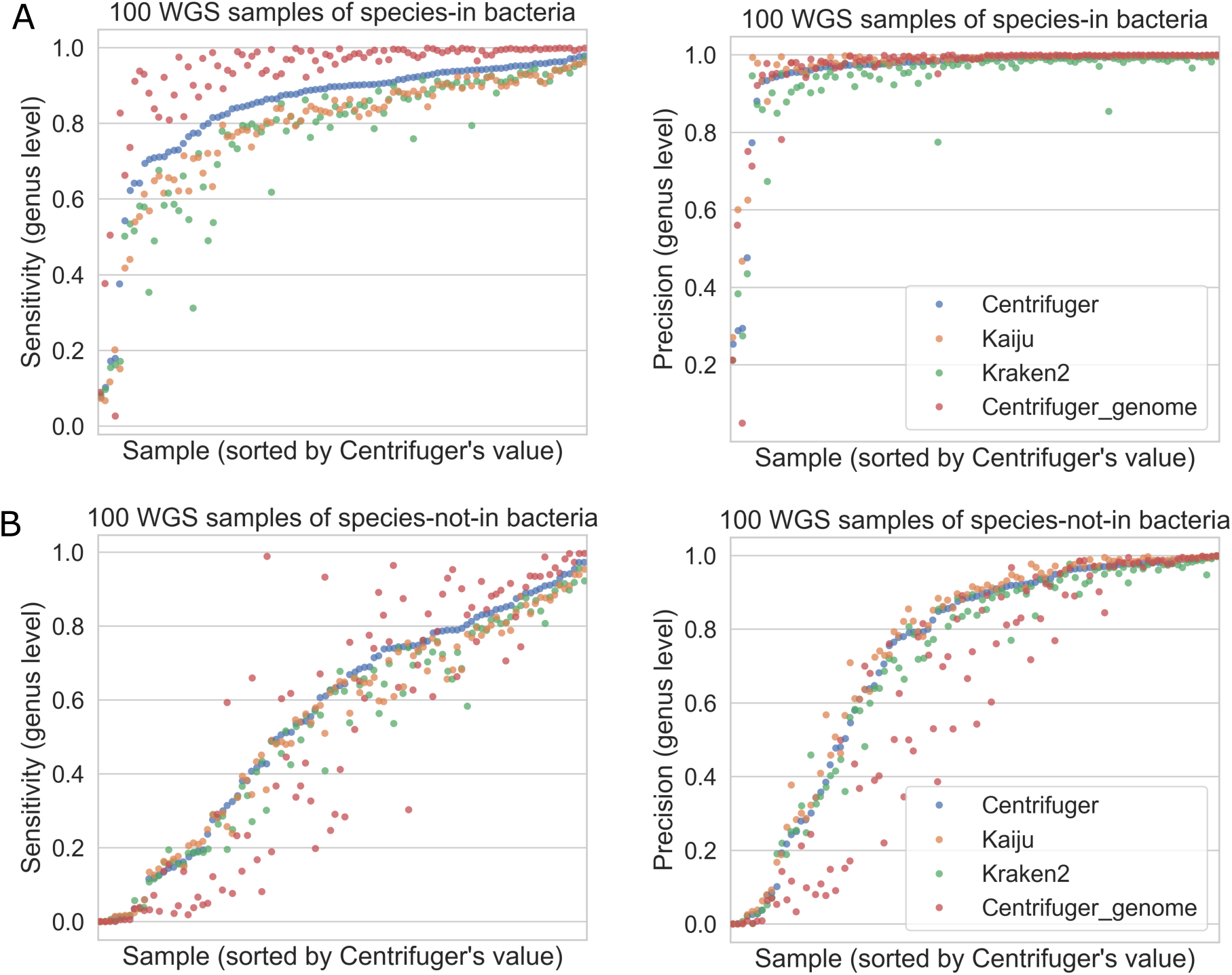
Performance of Centrifuger, Kaiju, and Kraken2 on bacterial WGS datasets (A) Sensitivity (left) and precision (right) if species of bacteria are present in the database. (B) Sensitivity (left) and precision (right) if species of bacteria are not in the database but their genera are present in the database.

For the species-not-in scenario, Centrifuger was 4.8% more sensitive than Kaiju, but was 2.5% less precise (**Figure 5B**), on average. Therefore, Centrifuger had a slightly higher mean F1 score than Kaiju’s (0.623 vs 0.611, **Figure S2**). Centrifuger outperformed Kraken2, with the mean sensitivity 7.1% higher and the mean precision 2.9% higher. In this scenario, Centrifuger_genome was substantially worse than the protein-based classifications, supporting that searching the protein database can alleviate the database incompleteness issue.

### SARS-CoV-2 scRNA-seq data

ScRNA-seq allows us to inspect the transcriptome at cellular resolution, and it also captures the microbial RNAs from a cell^27^. Centrifuger can parse the cell barcode information in scRNA-seq data, facilitating the study of microbial activity in different cell populations. We applied Centrifuger to the 10x Genomics scRNA-seq sample, SRR11181956^28^, from bronchoalveolar lavage fluid of a patient with severe SARS-CoV-2 infection to study the viral infection in different cell types. For human scRNA-seq data, we needed a database containing comprehensive host information because the human reference genome was often insufficient for filtering the host reads that may confound the microbiome analysis^29^.

Therefore, in this experiment, we created a Centrifuger index from the NCBI nr database, containing over 596 million protein sequences with a total of about 250 billion AAs. The average BWT sequence run length was 4.2 for nr protein sequences, and the Centrifuger index was 182 GB in size. The scRNA-seq data was processed with Cell Ranger v8.0.1, which generated an alignment BAM file. Centrifuger then conducted taxonomic classification on the unmapped reads against the nr database, and identified 2,294 reads uniquely classified to SARS-CoV-2 (taxonomy ID: 2697049).

To analyze in which cell types SARS-CoV-2 was most active, we first conducted conventional scRNA-seq data analysis, including quality control and clustering (**Figure 6A**, Methods). The SRR11181956 sample was part of the Viral-Track^27^ study that utilized mappings on the virus reference genomes to study the viral transcriptome in scRNA-seq data. This study included the cell type annotation, so we followed their annotation resolution and grouped the cells into three types: lymphocytes, myeloid cells, and epithelial cell (**Figure 6B, Figure S3**). We observed that some clusters of myeloid and epithelial cells had substantially higher fractions of cells associated with SARS-CoV-2 reads identified by Centrifuger (**Figure 6C**). In particular, the myeloid cells that were enriched with SARS-CoV-2 reads corresponded to the SPP1^hi^C1QA^hi^ macrophages (**Figure 6D**), which was consistent with the Viral-Track study.

**Figure 6.**
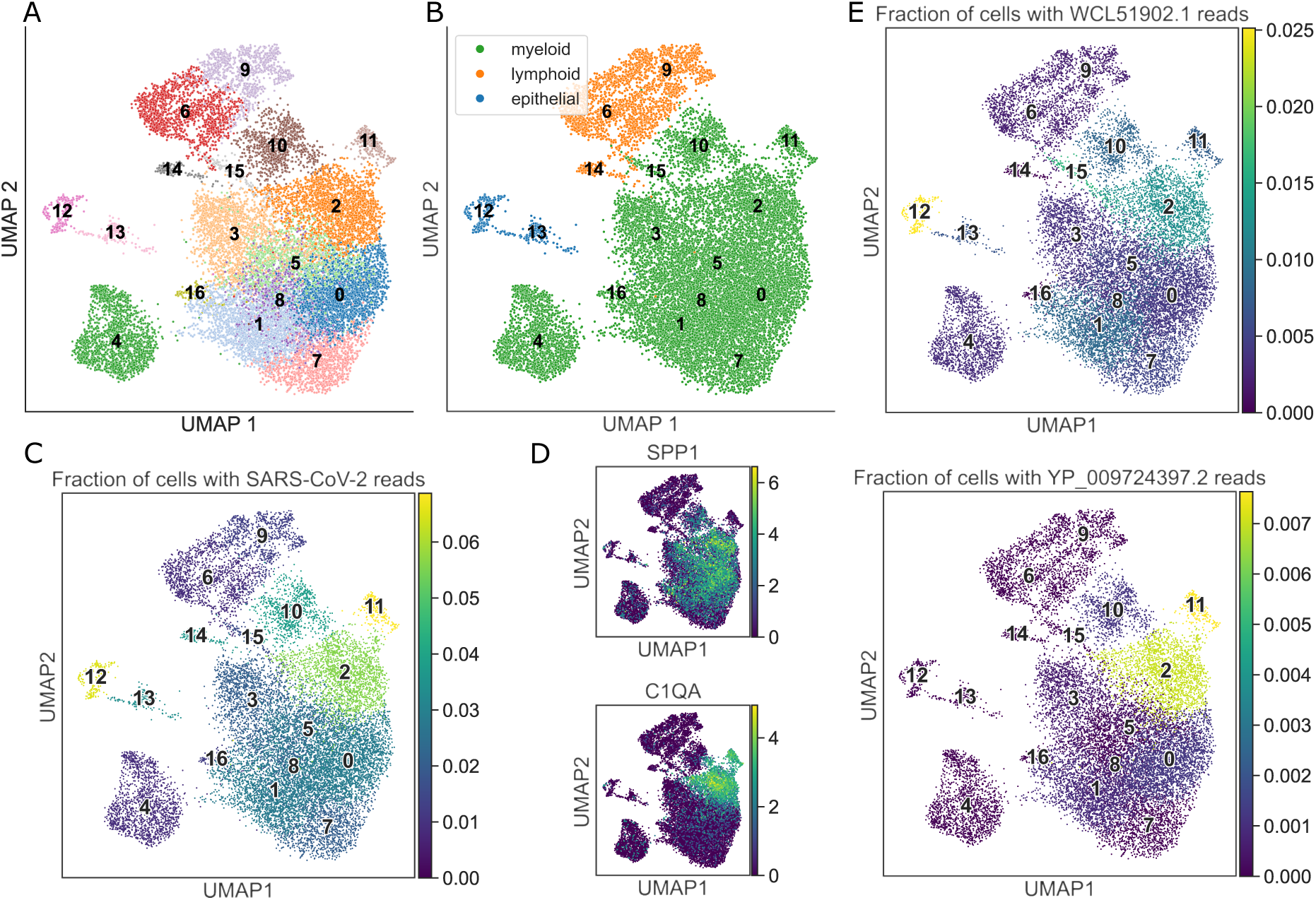
Analysis of the scRNA-seq data from a sample with severe SARS-CoV-2 infection. (A) UMAP and clusters of the SRR11181956 dataset. (B) Cell type annotation overlay on UMAP. (C) Fraction of cells with SARS-CoV-2 reads for each cluster. (D) The gene expression distribution of *SPP1* and *C1QA* genes. (E) Fraction of cells with reads classified to WCL51902 (ORF1ab polyprotein, top) and YP_009724397 (nucleocapsid phosphoprotein, bottom) for each cluster.

The sequence ID information in the index allows Centrifuger to find the protein ID for the reads. The two proteins with the most classified reads were WCL51902 with 654 reads and YP_009724397 with 650 reads. WCL51902 corresponds to the ORF1ab polyprotein, which spans a long genome region, and YP_009724397 corresponds to the SARS-CoV-2 nucleocapsid phosphoprotein (N protein) that is encoded by the genomic sequence on the 3’ end. This was consistent with the pseudo-bulk read alignment coverage analysis on the SARS-CoV-2 genome in the Viral-Track study. We observed that epithelial cells had a higher fraction of cells with the WCL51902 reads, while SPP1^hi^C1QA^hi^ macrophages were relatively enriched with the YP_009724397 reads, suggesting that SARS-CoV-2 may have different transcription patterns or infection effects in the two cell types (**Figure 6E**). Specifically, our data reflected that SARS-CoV-2 mainly infected alveolar epithelial cells^30^. Since C1Q is a marker for macrophage phagocytosis^31,32^, our finding suggested that SARS-CoV-2 could still be active after phagocytosis and started to upregulate N protein gene expression. Although not observed in this dataset, the SARS-CoV-2 N proteins in macrophages could induce a cytokine storm^33^.

## Discussion

In the actual implementation, we utilize the wavelet tree in the shape of complete binary tree to represent RBBWT-1S because its rank query time and space usage are affected by the alphabet size by a factor of *O*(log *σ*). This also allows the data structure to efficiently handle a variety of text types.

Nevertheless, for Centrifuger’s application, the alphabet set is known. Therefore, we can improve Centrifuger’s efficiency by incorporating more efficient representations that are optimized for nucleotide and AA alphabets, such as AWFM-index^34^ and flattened bit vectors^35^.

In this work, we show that searching a protein database alleviates the issue of database incompleteness and provides taxonomic information for reads from species missing in the database. However, this is at the expense of classification specificity compared to using the genome database, especially when the species genome is known. Metabuli has proposed the search paradigm through hybrid k-mers, i.e., metamer, where a k-mer contains translated AA sequence and original genomic sequence information. For classification, Metabuli can use the AA sequence portion to find candidate metamers, and then identify the metamer with the shortest Hamming distance. As a result, Metabuli can effectively utilize genomic information and find underrepresented species with high specificity. Future work is needed to explore the implementation of hybrid sequences for the FM-index to combine the advantages of searching genome databases and protein databases.

We applied Centrifuger to study the SARS-CoV-2 transcriptome in the scRNA-seq data from a severely infected patient. We identified that SARS-CoV-2 may express different transcripts in macrophages and epithelial cells by classifying reads against the NCBI nr database. However, the nr database has many highly similar protein sequences, so the classification results usually cannot be resolved at the sequence ID level. Therefore, this discovery was based on an additional Centrifuger run that exhaustively reports the sequence IDs to which a SARS-CoV-2 read was classified. One way to streamline the analysis in the future is to incorporate protein cluster information, such as UniRef90^36^ and ClusteredNR database, into the index to report the cluster ID if the sequence ID is unreachable.

In sum, we have designed a new lossless compressed data structure, RBBWT-1S, to reduce the size of the FM-index for sequences with relatively larger alphabet size, like protein sequences. We show that RBBWT-1S supports substantially faster rank queries than the original RBBWT at the expense of small space usage overhead for the AA alphabet. We implement RBBWT-1S in Centrifuger for taxonomic classification against protein databases like NCBI RefSeq and nr. In various benchmarks, Centrifuger shows higher sensitivity than commonly used methods Kaiju and Kraken2, with similar precision.

Centrifuger is substantially more memory-efficient while having the classification speed comparable to that of Kaiju, which utilizes a plain FM-index. Centrifuger not only finds taxonomy IDs, but also reports the sequence ID, providing additional valuable transcriptomic information for data like scRNA-seq data. Centrifuger is free open-source software under the MIT license and is available at: https://github.com/mourisl/centrifuger.

## Methods

### Sequencing data and benchmarks

We generated the simulated Illumina data using ART v2.5.8 from the 67,673 RefSeq bacteria, archaea, and virus complete genome sequences downloaded in August 2025. The options for ART were “art_illumina -ss HS25 -l 100 -m 1000 -s 100 -f 0.003”, which generated about 3.4 million read pairs. To form the final simulated data, we then randomly selected 1 million read pairs whose corresponding genomes had valid taxonomy information. For the simulated data from CAMI2, we downloaded the first 10 samples from the strain-madness dataset at https://frl.publisso.de/data/frl:6425521/strain/short_read/. For the bacterial WGS samples, we used the SraRunInfo file downloaded in our previous work^19^ with the search keyword “(“Bacteria”[Organism] OR “Bacteria Latreille et al. 1825”[Organism]) AND (“2022/01/01”[MDAT]: “2023/08/01”[MDAT]) AND (“biomol dna”[Properties] AND “strategy wgs”[Properties] AND “platform illumina”[Properties] AND “filetype fastq”[Properties])”. We re-selected 100 samples without duplicated species for the species-in scenario and 100 samples without duplicated genera for species-not-in scenario (**Table S2**). The selected samples had read fragment numbers between 1 million and 5 million to ensure the benchmark could be finished within a reasonable time. For the scRNA-seq data, we selected the sample SRR11181956, the largest patient scRNA-seq sample from the project PRJNA608742^28^. This sample was sequenced from poly-A-selected RNAs in the bronchoalveolar lavage fluid of a severe SARS-CoV-2 patient using the 10x Genomics 5’-kit.

In this study, we compared Centrifuger v1.1.4, Kaiju v1.10.1, and Kraken2 v2.1.6. The running command for each method is described in **Table S3**. To build the index, we used the script “centrifuger-download” from Centrifuger to download the taxonomy tree structure and the protein sequences for each of the 67,498 bacteria, archaea, and virus species with the complete genome from RefSeq. Since many species can produce the same protein, we deduplicated the protein sequences. If a protein ID, i.e., NCBI accession ID, was found in multiple taxa, we only kept one sequence copy and its taxonomy ID was the lowest common ancestor of these taxa. Centrifuger’s and Kaiju’s indexes were created on the deduplicated protein sequences. Kraken2 stored the minimizers and their taxa’s lowest common ancestor information in the index, so it did not need this deduplication step. We note that the number of genomes for generating the simulated data was slightly greater than the number of taxa for building the protein index, which could be due to the database inconsistencies or unannotated genomes.

All the benchmarks were conducted on a dedicated AMD EPYC 9354 processor machine with 768 GB of memory. The memory footprint was measured as the “Maximum resident set size” value from the “/usr/bin/time -v” command. When measuring speed, each classifier was run four times with a single thread on this dedicated server node. The reported classification speed was calculated by taking the fastest runtime after excluding index loading time.

### Run-block compression with one sequence

In run-block compression, a sequence *T*, such as the BWT sequence, is split into fixed-size blocks, denoted as *T*_1_, *T*_2_, … , *T_m_*, where *n* = |*T*|, 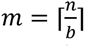 is the number of blocks, and *b* is the block size. Without loss of generality, we assume n is a multiple of b. A block *T_i_* is a run block if it has a single run of characters, e.g., *T_i_* = *c_i_^b^* for a character *c_i_*. So this run block can be represented with one character, thus reducing the space usage. We define the term 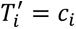 if it’s a run block, and 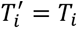 if not. In other words, *T′* represents the compressed form of a block *T_i_* depending on whether it is a run block or not. In the original run-block compression scheme, we create two sequences, corresponding to the *T′*^′^ from the run blocks and non-run blocks, respectively. In the run-block compression with one sequence scheme, we directly create one compressed sequence 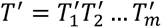. For example, for the raw sequence *T* =” LLLLLASGLLLL”, its *T*^′^ = “LLASGL”, with 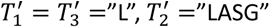 when *b* = 4. As in RBBWT, we have an additional bit vector *B_R_* to indicate whether *T_i_* is a run block or not. So in this example, *B_R_* = 101. In RBBWT-1S, we need a bit vector *B^C^_R_* for each character *c* ∈ Σ, indicating whether each of the *c* in *T*^′^ is from a run block or not. In the example, we have 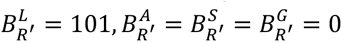 . Note that the total length of character-wise bit vectors equals the length of *T*^′^, i.e., 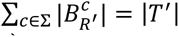 In the work of RBBWT^19^, we showed that |*B_R_*| and |*T*’| are on the order of 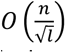 in the worst case, where *l* = *n*/*r* is the average run length. Therefore, the space usage of run-block compression with one sequence is also 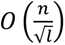 words and requires about 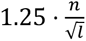 more bits than RBBWT due to storing *B^C^_R_* when using the rank9 method^37^ to support bit-vector rank queries.

The rank query, denoted as *rank_c_*(*i*, *T*), finds the number of occurrences of the character *c* in a sequence *T* before and on its position *i*, where we use a 1-based index in proofs and *rank_c_*(0, *T*) = 0. We can represent a sequence using a wavelet tree, which supports rank queries with the time complexity *O*(log *σ*) and the space complexity *O*(*n* log *σ*) bits.

**Theorem 1.** The time complexity for the rank query *rank_c_*(*i*, *T*) using run-block compression with one sequence can be *O*(log *σ*).

Proof: Let *k* = ⌈*i*/*b*⌉ denote the block that *i* resides in, and *r_R_* = *rank*_1_(*k* − 1, *B_R_*) represent the number of run blocks before block *k*. Since *i* can be inside a run block, we use the variable *x* = *I_BR_*[*_k_*]_=1_ ⋅ (*i* − 1)%*b* representing that *i* is *x* away from the block start, where *I* _∗_ is an indicator function that equals 1 if the subscript is true and 0 otherwise. With the notations, the position *i* of the original sequence *T* corresponds to the position *i*^′^ = *i* − *x* − (*b* − 1)*r_R_* in *T*’. Let 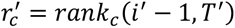 *r*^′^ = *rank* (*i*^′^ − 1, *T*^′^), then we can calculate the rank query on *T* by:

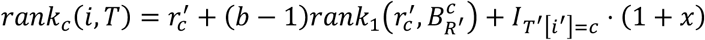

where the second term is to retrieve the count from the compressed run blocks of *c*. We deduct 1 from *i*′ in *r’_c_* so that the last term can properly handle the count if *T*^′^[*i*^′^] = *c*.

In practice, compared with the original run-block compression scheme, the new method replaced the cost of one rank query on the sequence by a rank query on the bit vector. Since we can use the wavelet tree to represent *T*’ and the rank query on a bit vector takes *O*(1) time, the overall time complexity is *O*(log *σ*). □

We note that the space complexity for representing *T*’ can be reduced to *O*M|*T*^′^| ⋅ *H*_0_(*T*′)N if the wavelet tree is Huffman-shaped^38^, where *H*_0_ is the Shannon entropy of the sequence. However, the actual taxonomic classification involves searching all possible reading frames of a read, and the translated AAs can have a very different frequency pattern from that of the indexed database. As a result, too many AA rank queries need to access the deep levels of the Huffman tree and slow down the classification process. Furthermore, the space savings can be small. For example, the *H*_0_(*T*^′^) for the RefSeq prokaryotic and viral protein sequences is about 4.14, where the maximum entropy for the 21 letters is about log_2_ *σ*=4.39. Therefore, we still use the complete binary tree shape for the wavelet tree.

### Index construction

For each protein sequence, Centrifuger appends a special character “$” to mark the sequence ending. This strategy is different from Centrifuger_genome that uses a hash table to track the end of a sequence. This is because genomic sequences are usually long, and the nucleotide alphabet size 4 has good computational properties. For example, the wavelet tree can be a full tree without wasting any bit information. The BWT sequence is constructed based on the sorted suffix array using the blockwise suffix sorting algorithm^39^, where “$” is lexicographically less than other characters. This procedure also generates the sampled suffix array elements as in the standard FM-index, and Centrifuger will convert the suffix array values to the corresponding sequence IDs to reduce the index size. There are two types of sampled suffix array values: the positions corresponding to “$” and the positions that are multiples of a user-specified value, 16 by default. The positions corresponding to “$” are at the beginning of the suffix array, so both sampled suffix array values can be stored in arrays and it is straightforward to know whether a suffix array position is sampled. Since the “$” marks the end of a sequence and the search is backward, when converting its suffix array value to the sequence ID, Centrifuger actually uses the next sequence’s ID.

To estimate the block size b, we split the BWT sequence into 1024 regions and create a sequence by concatenating the first 1024 AAs from each region. This creates a sequence with about 1 million AAs. Centrifuger then tests the block sizes that are powers of 2, and another value 1048576/√*l* after estimating its average run length *l*. The block size that leads to the highest compression ratio will be the final block size. In the previous Centrifuger version, we inferred the block size from the first 1 million characters of the BWT. However, in the protein database, the first 1 million positions of the BWT sequence correspond to the characters before the “$” symbol, i.e., the end of the protein. Therefore, the previous Centrifuger strategy would create inaccurate run block size estimation for the full BWT sequence from a protein database.

### Taxonomic classification

Centrifuger greedily searches for semi-maximal exact matches on the translated AA sequences, where a match is semi-maximal if it cannot be extended further backward. For a read strand, Centrifuger will translate it into three AA sequences, each corresponding to a reading frame. Therefore, a read will be translated into six AA sequences. For each sequence, Centrifuger starts the backward search from its end to find the first semi-maximal exact match. Centrifuger then skips one AA and starts a new backward search to locate the next semi-maximal exact match. For the translated AA sequence, the stop codon is represented as “_” and the codon containing the uncertain base “N” in the raw read will be translated to “?”. These special characters will immediately trigger a mismatch in the backward search.

The next step is to retrieve the taxonomy ID from selected matches. Centrifuger will first filter short matches that are likely to occur by chance. Specifically, for a match *M* of length *l_M_*, we filter it if *l_M_* < 11 or 2*n*/21*^lM^* > 0.01 by default. Centrifuger then assigns the score (*l_M_* − 5)^2^ to the match *M*. This scoring function is an empirical function based on Centrifuger_genome, where we modify the offset from 15 to 5 because a codon corresponds to three nucleotides. Then, Centrifuger picks the reading frame with the highest sum of scores from its matches among the six possible reading frames. In the example of Figure 1, reading frame 3 achieves the highest score. For the paired-end data, Centrifuger picks the highest-scoring pair of reading frames from different strands for the two reads. For each valid match of the selected reading frame, Centrifuger applies LF-mapping to locate the sampled sequence ID information. Specifically, a match *M* corresponds to the position interval [*s_M_*, *e_M_*] on the BWT. As in a conventional FM-index, we can apply the LF mapping for each element in this interval until reaching the suffix position with the sampled sequence ID. We then add the score (*l_M_* − 5)^2^ to that sequence ID. If the match *M* is found more than once in a reference protein sequence, Centrifuger only records the score once. In the end, the highest-scoring sequence IDs are the classification result. In the example of Figure 1, sequence ID “X” is found in two matches of length 20 and 11, respectively, so it has the highest score with the value of (20 − 5)^2^ + (11 − 5)^2^ = 261. In addition to sequence IDs, Centrifuger reports their corresponding taxonomy IDs.

When the number of highest-scoring sequence IDs for a query is above the per-read report limit, denoted by k (default k=1), Centrifuger will reduce the output entries by moving up the taxonomy IDs along the taxonomy tree until the number of distinct taxa is within k. So by default, Centrifuger reports the taxonomy ID of the lowest common ancestor for each read fragment. The value k affects how sequence IDs are resolved for a match. If *e_M_* − *s_M_* + 1 > 40 ⋅ *k*, we will only select 40 ⋅ *k* elements that are evenly distributed in the interval [*s_M_*, *e_M_*] to get the sequence IDs. Although resolving only a subset of sequence IDs may miss the true ID, this strategy aims to find the right taxonomy ID at higher taxonomic ranks. The factor 40 is user-adjustable. When k=0, Centrifuger will exhaustively report all the highest-scoring sequence IDs. All these approaches are based on Centrifuger_genome.

### SARS-CoV-2 scRNA-seq data processing

We followed common practice^40^ to analyze the scRNA-seq data using Scanpy^41^ v1.12. We first conducted quality control by filtering cells whose log(1+x) values of the total number of reads and the total number of detected genes were outside 5 times the median absolute deviation (MAD). Cells with the percentage of mitochondrial reads above the value of 8 or outside 3 times the MAD were filtered. We also filtered genes that were detected in fewer than 20 cells. The filtering gave a count matrix with 17,543 cells and 20,410 genes. Scrublet^42^ did not detect any doublets in this unnormalized count matrix using the parameter expected_doublet_rate=0.12 (default 0.05), a value based on the guideline for 15,000 cells. The gene expression values for each cell were then normalized and transformed using the log1p function. The top 2,000 highly variable genes were retained for the principal component analysis (PCA), and the top 50 principal components were utilized for generating the neighborhood graph. The UMAP^43^ coordinates and cluster information from the Leiden method^44^ were based on this neighborhood graph. To facilitate the cell type annotation, we applied the method CellTypist^45^ v1.7.1 for automated cell type annotation using the “COVID19_Immune_Landscape” model as the reference followed by cluster-based majority voting (**Figure S3**).

For the nr database, the protein sequences were extracted through “blastdbcmd” on the BLAST database downloaded in January 2026. The sequence ID to taxonomy ID mapping information was based on the prot.accession2taxid file from https://ftp.ncbi.nlm.nih.gov/pub/taxonomy/accession2taxid/. Although the nr database was non-redundant, there were many highly similar protein sequences. For example, proteins YP_009724397 and WCA99607 differ only by one AA, where the second AA is “S” in YP_009724397 and “Y” in WCA99607. As a result, Centrifuger only yielded sequence IDs for 632 out of the 2,294 SARS-CoV-2 reads. Therefore, we extracted the reads classified to the taxonomy ID 2697049 and reran Centrifuger with the option “-k 0” to report all the highest-scoring protein sequences for each read. We attempted to create the full nr index for Kaiju, but the job was killed due to the out_of_memory error on our server.

## Data and code availability

The source code of Centrifuger is available at https://github.com/mourisl/centrifuger. Centrifuger v1.1.4 was used in this study. The code for the evaluations and experiments is available at https://github.com/mourisl/centrifuger_protein_evaluations. The NCBI SRA accession numbers of the WGS datasets are listed in Table S2. The Centrifuger index for the RefSeq prokaryotic and viral protein sequences is available at Zenodo doi:10.5281/zenodo.22663514. The nr index is publicly accessible on Dropbox as described in the Centrifuger documentation and can be downloaded through “centrifuger-download”.

## Acknowledgements

This work is supported by the National Institute of Allergy and Infectious Diseases of the National Institutes of Health under grant number R01AI195878 (L.S.) and the National Institute of General Medical Sciences of NIH under grant number P20GM130454 (Dartmouth). One hundred percent of the total costs of this project (about $200,000) is financed from federal grants. The content is solely the responsibility of the authors and does not necessarily represent the official views of the National Institutes of Health.

## Authors’ contributions

L.S. conceived the project, designed the algorithm, implemented and evaluated the software, and wrote the manuscript.

## Supplementary Figures

**Figure S1.**
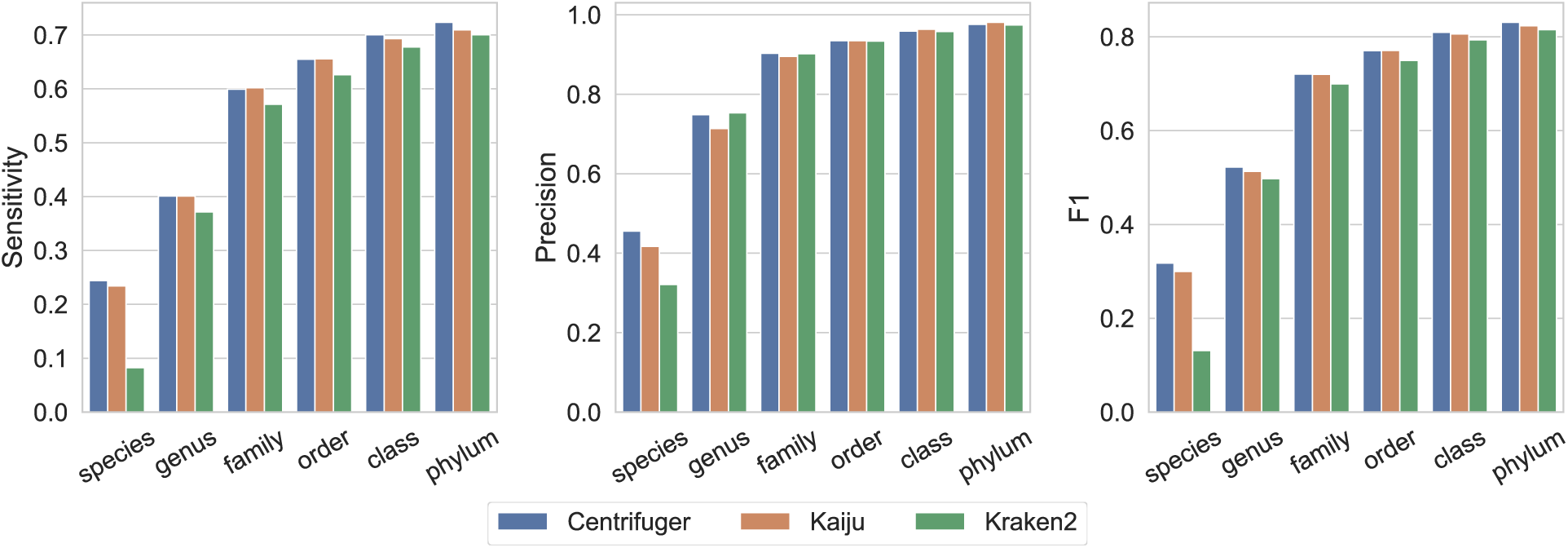
Performance of Centrifuger, Kaiju, and Kraken2 on classifying the simulated reads against a trimmed database with one representative strain per genus. The reads from those representatives are removed.

**Figure S2.**
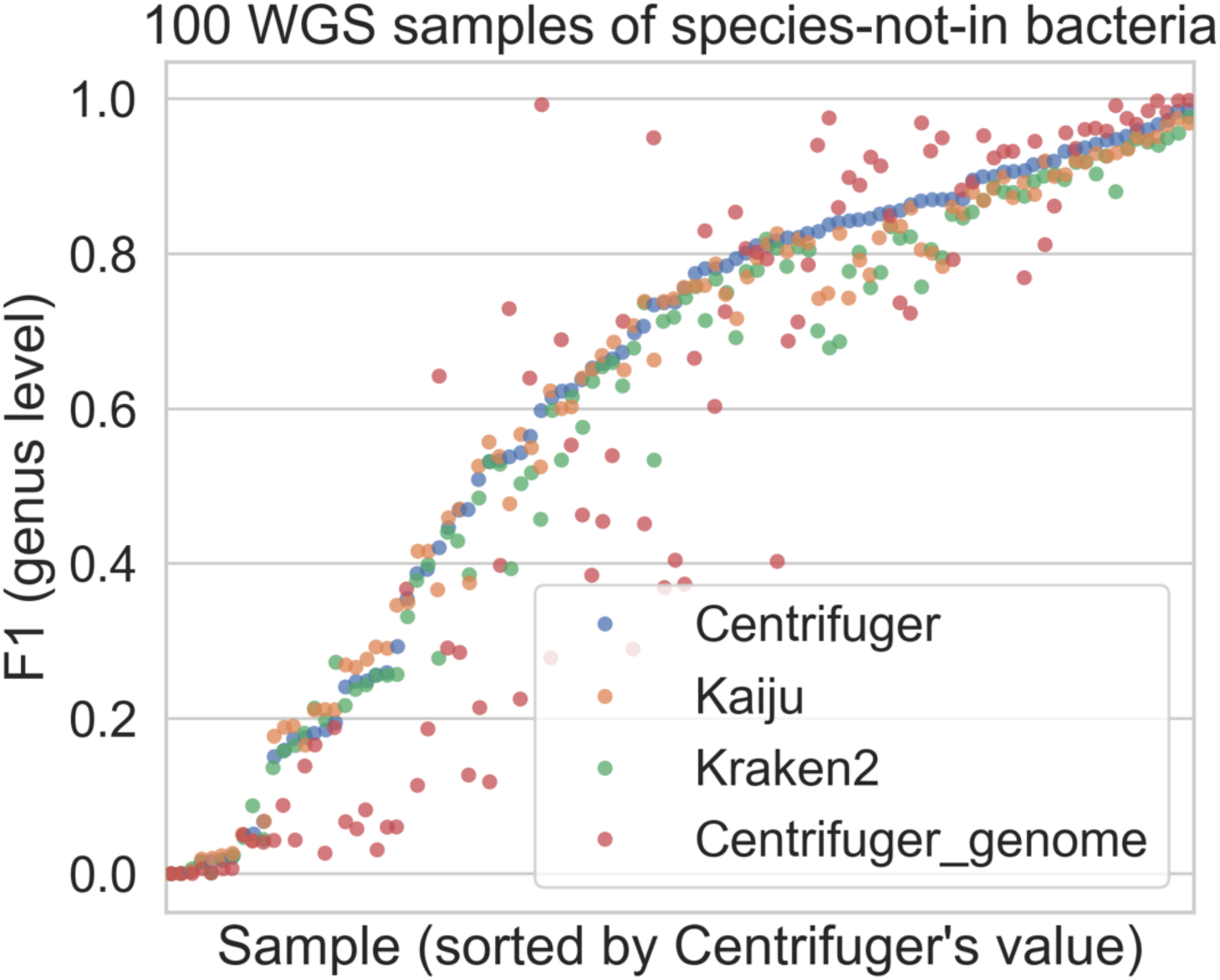
F1 scores of Centrifuger, Kaiju and Kraken2 on bacterial WGS datasets when the species is missing in the reference database but their genera are present

**Figure S3.**
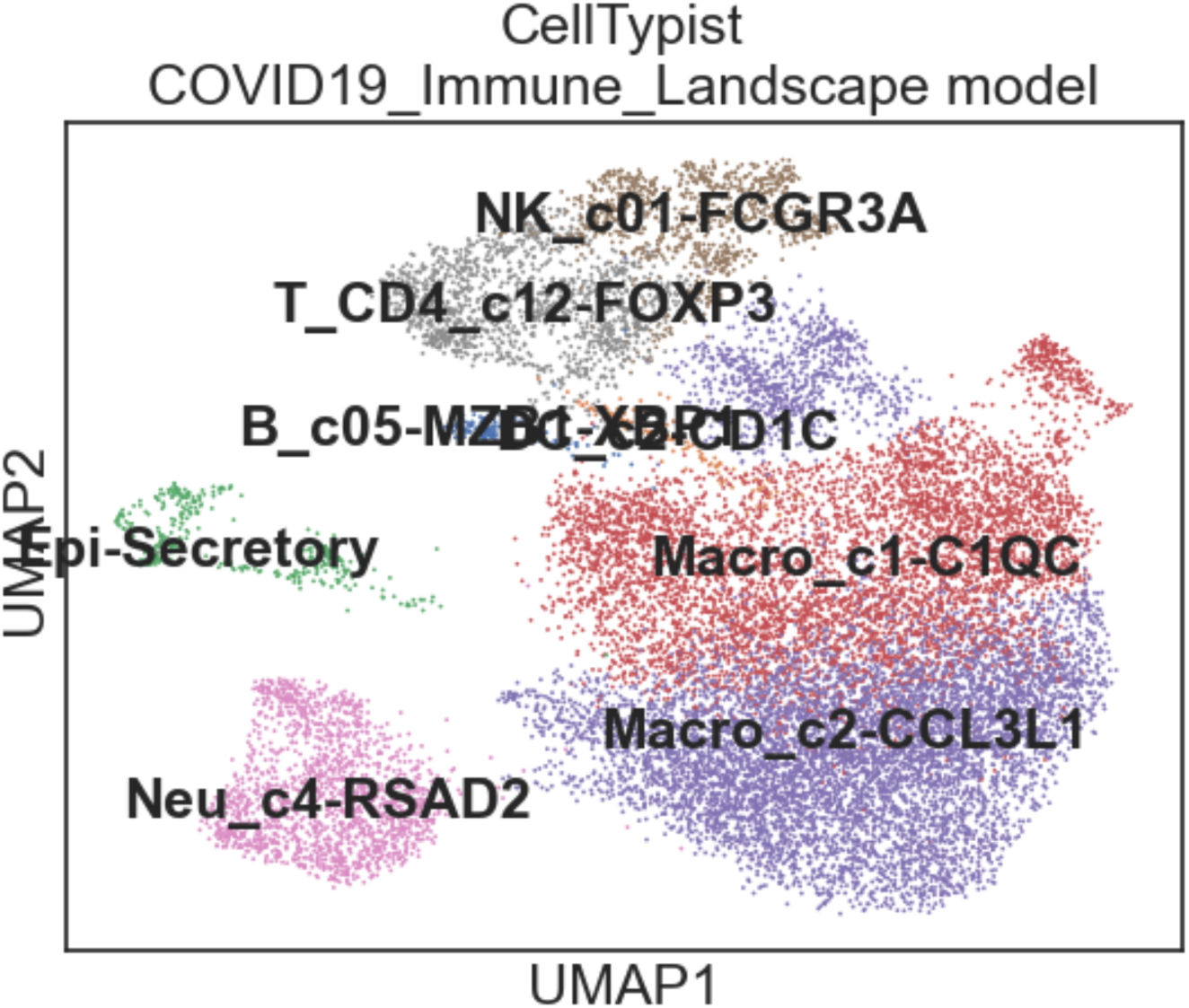
Automated cell type annotation from CellTypist for the SARS-CoV-2 scRNA-seq data analysis

## Supplementary Tables

**Table S1.** The classification accuracy across taxonomic ranks in the ART-generated simulated data Sensitivity=TP/T, Precision=TP/P. The highest values of sensitivity and precision at each rank are bolded. The reduced number of cases, i.e., the number of simulated read fragments, in higher taxonomic ranks is due to the missing ranks in the taxonomy tree.

| rank | matched (TP) | predicted (P) | cases (T) | sensitivity | precision | method |
| --- | --- | --- | --- | --- | --- | --- |
| strain | 147629 | 167731 | 1000000 | **0.1476** | **0.8802** | Centrifuger |
| strain | 137300 | 188677 | 1000000 | 0.1373 | 0.7277 | Kaiju |
| strain | 123215 | 155607 | 1000000 | 0.1232 | 0.7918 | Kraken2 |
| species | 330308 | 340947 | 1000000 | **0.3303** | 0.9688 | Centrifuger |
| species | 283812 | 291067 | 1000000 | 0.2838 | **0.9751** | Kaiju |
| species | 268610 | 293079 | 1000000 | 0.2686 | 0.9165 | Kraken2 |
| genus | 787074 | 792296 | 999726 | **0.7873** | 0.9934 | Centrifuger |
| genus | 740586 | 744975 | 999726 | 0.7408 | **0.9941** | Kaiju |
| genus | 755455 | 771350 | 999726 | 0.7557 | 0.9794 | Kraken2 |
| family | 923043 | 924748 | 997184 | **0.9256** | **0.9982** | Centrifuger |
| family | 887721 | 889685 | 997184 | 0.8902 | 0.9978 | Kaiju |
| family | 884763 | 892486 | 997184 | 0.8873 | 0.9913 | Kraken2 |
| order | 953132 | 954301 | 997732 | **0.9553** | **0.9988** | Centrifuger |
| order | 919124 | 920274 | 997732 | 0.9212 | **0.9988** | Kaiju |
| order | 918217 | 924503 | 997732 | 0.9203 | 0.9932 | Kraken2 |
| class | 962271 | 962974 | 993614 | **0.9685** | **0.9993** | Centrifuger |
| class | 929145 | 929784 | 993614 | 0.9351 | **0.9993** | Kaiju |
| class | 944673 | 949073 | 993614 | 0.9507 | 0.9954 | Kraken2 |
| phylum | 972847 | 973382 | 999746 | **0.9731** | 0.9995 | Centrifuger |
| phylum | 939482 | 939897 | 999746 | 0.9397 | **0.9996** | Kaiju |
| phylum | 958204 | 961453 | 999746 | 0.9584 | 0.9966 | Kraken2 |

**Table S2.**
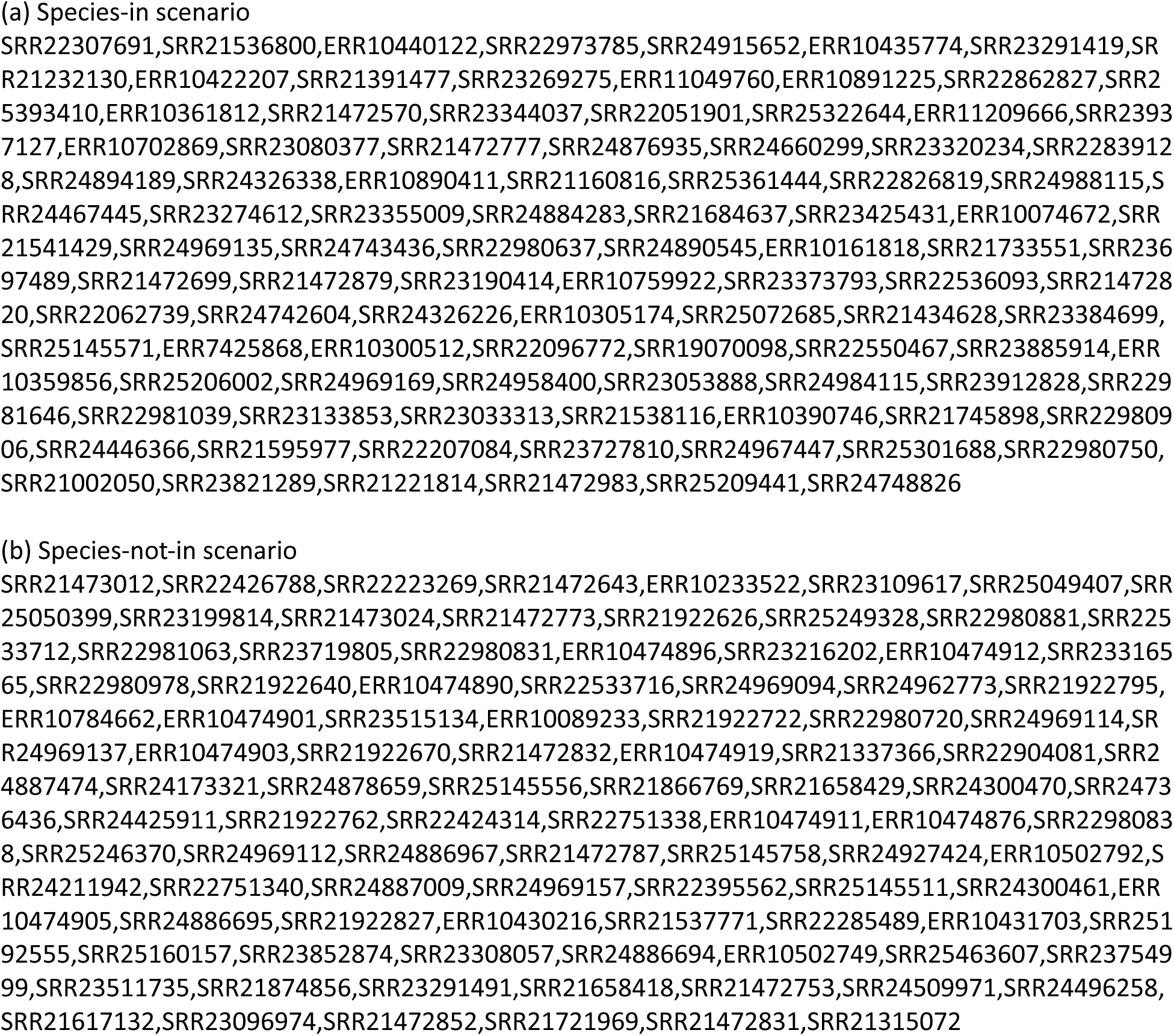
SRA IDs for the samples used in the bacterial WGS classification evaluations

**Table S3.**
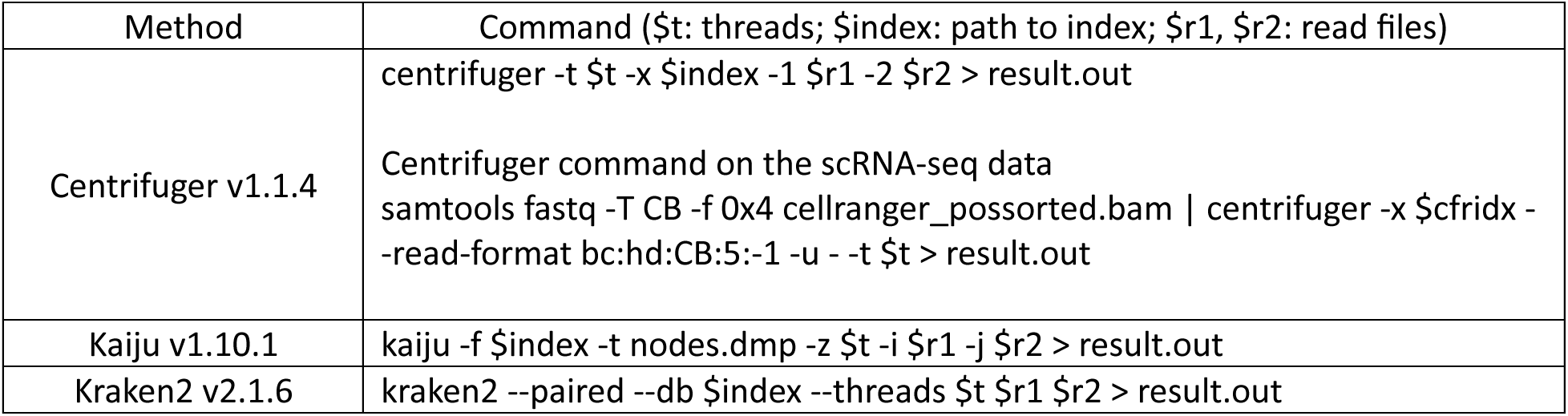
Running commands for the classifiers used in the evaluations

## References

1. Tringe, S. G. & Rubin, E. M. Metagenomics: DNA sequencing of environmental samples. Nat Rev Genet 6, 805–814 (2005).

2. Zhang, L. et al. Advances in Metagenomics and Its Application in Environmental Microorganisms. Frontiers in Microbiology 12, (2021).

3. Chiu, C. Y. & Miller, S. A. Clinical metagenomics. Nat Rev Genet 20, 341–355 (2019).

4. Knight, R. et al. Best practices for analysing microbiomes. Nat Rev Microbiol 16, 410–422 (2018).

5. Sirasani, J. P., Gardner, C., Jung, G., Lee, H. & Ahn, T.-H. Bioinformatic approaches to blood and tissue microbiome analyses: challenges and perspectives. Brief Bioinform 26, bbaf176 (2025).

6. Pruitt, K. D., Tatusova, T. & Maglott, D. R. NCBI reference sequences (RefSeq): a curated non-redundant sequence database of genomes, transcripts and proteins. Nucleic Acids Res 35, D61–D65 (2007).

7. Benson, D. A. et al. GenBank. Nucleic Acids Res 41, D36–42 (2013).

8. Parks, D. H. et al. GTDB: an ongoing census of bacterial and archaeal diversity through a phylogenetically consistent, rank normalized and complete genome-based taxonomy. Nucleic Acids Research 50, D785–D794 (2022).

9. Wood, D. E., Lu, J. & Langmead, B. Improved metagenomic analysis with Kraken 2. Genome Biol 20, 257 (2019).

10. Ounit, R., Wanamaker, S., Close, T. J. & Lonardi, S. CLARK: fast and accurate classification of metagenomic and genomic sequences using discriminative k-mers. BMC Genomics 16, 236 (2015).

11. Shen, W. et al. KMCP: accurate metagenomic profiling of both prokaryotic and viral populations by pseudo-mapping. Bioinformatics 39, btac845 (2023).

12. Piro, V. C. & Reinert, K. ganon2: up-to-date and scalable metagenomics analysis. NAR Genom Bioinform 7, lqaf094 (2025).

13. Kim, J. & Steinegger, M. Metabuli: sensitive and specific metagenomic classification via joint analysis of amino acid and DNA. Nat Methods 21, 971–973 (2024).

14. Nasko, D. J., Koren, S., Phillippy, A. M. & Treangen, T. J. RefSeq database growth influences the accuracy of k-mer-based lowest common ancestor species identification. Genome Biology 19, 165 (2018).

15. Commichaux, S., Luan, T., Muralidharan, H. S. & Pop, M. Database size positively correlates with the loss of species-level taxonomic resolution for the 16S rRNA and other prokaryotic marker genes. PLOS Computational Biology 20, e1012343 (2024).

16. Ferragina, P. & Manzini, G. Opportunistic data structures with applications. in Proceedings 41st Annual Symposium on Foundations of Computer Science 390–398 (2000). doi:10.1109/SFCS.2000.892127.

17. Kim, D., Song, L., Breitwieser, F. P. & Salzberg, S. L. Centrifuge: rapid and sensitive classification of metagenomic sequences. Genome Res 26, 1721–1729 (2016).

18. Menzel, P., Ng, K. L. & Krogh, A. Fast and sensitive taxonomic classification for metagenomics with Kaiju. Nat Commun 7, 11257 (2016).

19. Song, L. & Langmead, B. Centrifuger: lossless compression of microbial genomes for efficient and accurate metagenomic sequence classification. Genome Biol 25, 106 (2024).

20. Kreft, S. & Navarro, G. On compressing and indexing repetitive sequences. Theoretical Computer Science 483, 115–133 (2013).

21. Gagie, T., Gawrychowski, P., Kärkkäinen, J., Nekrich, Y. & Puglisi, S. J. A Faster Grammar-Based Self-Index. Preprint at 10.48550/arXiv.1109.3954 (2012).

22. Gagie, T., Navarro, G. & Prezza, N. Optimal-Time Text Indexing in BWT-runs Bounded Space. Preprint at 10.48550/arXiv.1705.10382 (2017).

23. Nishimoto, T. & Tabei, Y. Optimal-Time Queries on BWT-runs Compressed Indexes. Preprint at 10.48550/arXiv.2006.05104 (2021).

24. Mäkinen, V., Navarro, G., Sirén, J. & Välimäki, N. Storage and Retrieval of Highly Repetitive Sequence Collections. Journal of Computational Biology 17, 281–308 (2010).

25. Huang, W., Li, L., Myers, J. R. & Marth, G. T. ART: a next-generation sequencing read simulator. Bioinformatics 28, 593–594 (2012).

26. Meyer, F. et al. Critical Assessment of Metagenome Interpretation: the second round of challenges. Nat Methods 19, 429–440 (2022).

27. Bost, P. et al. Host-Viral Infection Maps Reveal Signatures of Severe COVID-19 Patients. Cell 181, 1475–1488.e12 (2020).

28. Liao, M. et al. Single-cell landscape of bronchoalveolar immune cells in patients with COVID-19. Nat Med 26, 842–844 (2020).

29. Gihawi, A. et al. Major data analysis errors invalidate cancer microbiome findings. mBio 0, e01607–23 (2023).

30. Sherman, A. C. et al. Acute SARS-CoV-2 infection. Nat Rev Dis Primers 11, 75 (2025).

31. Pulanco, M. C. et al. Complement protein C1q enhances macrophage foam cell survival and efferocytosis. J Immunol 198, 472–480 (2017).

32. Coulton, A. et al. Using a pan-cancer atlas to investigate tumour associated macrophages as regulators of immunotherapy response. Nat Commun 15, 5665 (2024).

33. Yao, Z. et al. SARS-CoV-2 nucleocapsid induces hyperinflammation and vascular leakage through the Toll-like receptor signaling axis in macrophages. Science Advances 12, eaea2780 (2026).

34. Anderson, T. & Wheeler, T. J. An optimized FM-index library for nucleotide and amino acid search. Algorithms for Molecular Biology 16, 25 (2021).

35. Gottlieb, S. G. & Reinert, K. Engineering rank queries on bit vectors and strings. Algorithms Mol Biol 20, 21 (2025).

36. The UniProt Consortium. UniProt: the Universal Protein Knowledgebase in 2025. Nucleic Acids Res 53, D609–D617 (2025).

37. Vigna, S. Broadword implementation of rank/select queries. in Proceedings of the 7th international conference on Experimental algorithms 154–168 (Springer-Verlag, Berlin, Heidelberg, 2008).

38. Barbay, J. & Navarro, G. On compressing permutations and adaptive sorting. Theoretical Computer Science 513, 109–123 (2013).

39. Kärkkäinen, J. Fast BWT in small space by blockwise suffix sorting. Theoretical Computer Science 387, 249–257 (2007).

40. Heumos, L. et al. Best practices for single-cell analysis across modalities. Nat Rev Genet 24, 550– 572 (2023).

41. Wolf, F. A., Angerer, P. & Theis, F. J. SCANPY: large-scale single-cell gene expression data analysis. Genome Biol 19, 15 (2018).

42. Wolock, S. L., Lopez, R. & Klein, A. M. Scrublet: Computational Identification of Cell Doublets in Single-Cell Transcriptomic Data. Cell Systems 8, 281–291.e9 (2019).

43. McInnes, L., Healy, J. & Melville, J. UMAP: Uniform Manifold Approximation and Projection for Dimension Reduction. Preprint at 10.48550/arXiv.1802.03426 (2020).

44. Traag, V. A., Waltman, L. & van Eck, N. J. From Louvain to Leiden: guaranteeing well-connected communities. Sci Rep 9, 5233 (2019).

45. Domínguez Conde, C., et al. Cross-tissue immune cell analysis reveals tissue-specific features in humans. Science 376, eabl5197 (2022).

